# Imaging large fields-of-view at high resolution in cryo-ET with square beam montaging

**DOI:** 10.64898/2026.08.27.747605

**Authors:** Eugene Y.D. Chua, Hamidreza Rahmani, James Zhen, Fabian Eisenstein, Yong Hyun Song, Jake D. Johnston, Hanyu Wang, Lambertus M. Alink, Mykhailo Kopylov, Chi-Min Ho, Danielle A Grotjahn, Alex de Marco

## Abstract

Visualizing macromolecules within their native cellular context by cryo-electron tomography (cryo-ET) is fundamentally limited by the trade-off between field of view and resolution: Capturing high-resolution information about biomolecules requires high magnification, which restricts the field of view and obscures the cellular context in which those biomolecules function. Collecting montage data by tiling the electron beam over the region of interest offers one solution, although traditional round electron beams cause excessive radiation damage across overlapping regions. We previously made electron beams square in shape, enabling montage collection with minimal overlap and thereby reducing excessive exposure and loss of high-resolution information. Here, we create a pipeline for collecting and processing montage cryo-ET data with square electron beams. We show that square beam montages retain high-resolution information by reconstructing virus-like particles to 3.5 Å resolution using sub-tomogram averaging, and apply the workflow to imaging a glial cell and malaria parasite lamellae over fields of view up to 65 µm^2^. We also provide a comprehensive protocol to make square beams accessible to the community.

## Introduction

Cryo-electron tomography (cryo-ET) is a powerful tool for visualizing the structure and organization of biomolecules in their native cellular environments at the nanometer scale (Anton et al., 2025; Chang et al., 2025; Klumpe et al., 2025; Mahamid et al., 2016; Schiøtz et al., 2024; Tegunov et al., 2021). For cryo-ET, biological samples are captured in a near-native frozen-hydrated state by flash freezing, then imaged in a cryo-transmission electron microscope (cryo-TEM) at different tilts to reconstruct a 3D volume (i.e., tomogram). To capture high-resolution information about biomolecules and study their function, high magnification (i.e., a small pixel size) is needed; however, this limits the field of view, obscuring the cellular context. Recent developments underscore the community’s interest in situating biological processes within their native contexts, including capturing low-magnification overviews to provide contextual information for high-magnification tomograms of some areas (Gebauer et al., 2026; Watson et al., 2026), and utilizing high-resolution 2D template matching to map the location and orientation of ribosomes in an entire lamella (Elferich et al., 2022).

Montage imaging, a method that tiles the electron beam across the region of interest to generate a montage, can also expand the field of view while imaging at high magnifications (Peck et al., 2022; Yang et al., 2023). Modern state-of-the-art cryo-TEMs come equipped with standard apertures that produce round beams that cannot be tiled without overlap. This is a challenge for frozen-hydrated biological specimens because they are damaged by the electron beam during imaging, and overlapping exposure regions receive excessive damage. Existing collection schemes aim to account for this by distributing the electron dose as evenly as possible across the field of view (Peck et al., 2022; Yang et al., 2023). However, the accumulated radiation damage from montaging with a round beam has been shown to preclude high-resolution sub-tomogram averaging reconstructions (Hylton et al., 2026).

To overcome the problem of accumulated radiation damage in overlapping montage regions, we developed a method to make electron beams square in shape (Brown et al., 2024; Chua et al., 2024). Such a beam can be tiled with minimal overlap, thereby avoiding excessive electron dose throughout the field of view and minimizing sample damage outside of the field-of-view. This initial method was sufficient to obtain montages that could be reconstructed into tomograms, in which particles such as ribosomes and apoferritin could be visualized (Chua et al., 2024). However, this technology has yet to be applied to sub-tomogram averaging and high-resolution imaging of cellular samples. In this work, we demonstrate the utility of square beam montages for cryo-ET applications by obtaining a 3.5 Å reconstruction of virus-like particles (VLPs) by sub-tomogram averaging (STA) from square montage tomograms. We then apply square montages to image a glial cell, enabling us to identify the polarity of microtubules found within the montage. Furthermore, we imaged lamellae of malaria parasites within red blood cells with fields-of-view up to 65 µm^2^, thereby maximizing both high-resolution and contextual information gathered from lamellae.

We also present an updated data collection strategy that uses minimal overlap between square tiles to align and stitch the montages, reducing tiling and stitching artifacts. We developed a custom data pre-processing workflow, including cropping unilluminated sensor areas and assembling the montage tilt series. We present a pipeline for processing square montage data, and accompanying detailed protocols for square aperture installation, software setup, data collection, and data processing. Our data processing pipeline utilizes tools commonly used by the cryo-ET community, allowing it to fit easily within existing workflows. Our goal is to increase accessibility for scientists who want to use the square aperture for montage tomography to investigate their biological system of interest.

## Results and Discussion

### A workffow for square montage data collection and processing

For square beam montage data collection, we updated PACE-tomo (Eisenstein et al., 2023) so that each target can be collected as a montage. PACE-tomo v1.9.3 now includes options to apply defocus gradient compensation between montage tiles, define square or rectangular montage shapes (e.g. 3×3, 3×5, 7×5, etc), and define the amount of overlap between tiles (Figure 1A).

**Figure 1.**
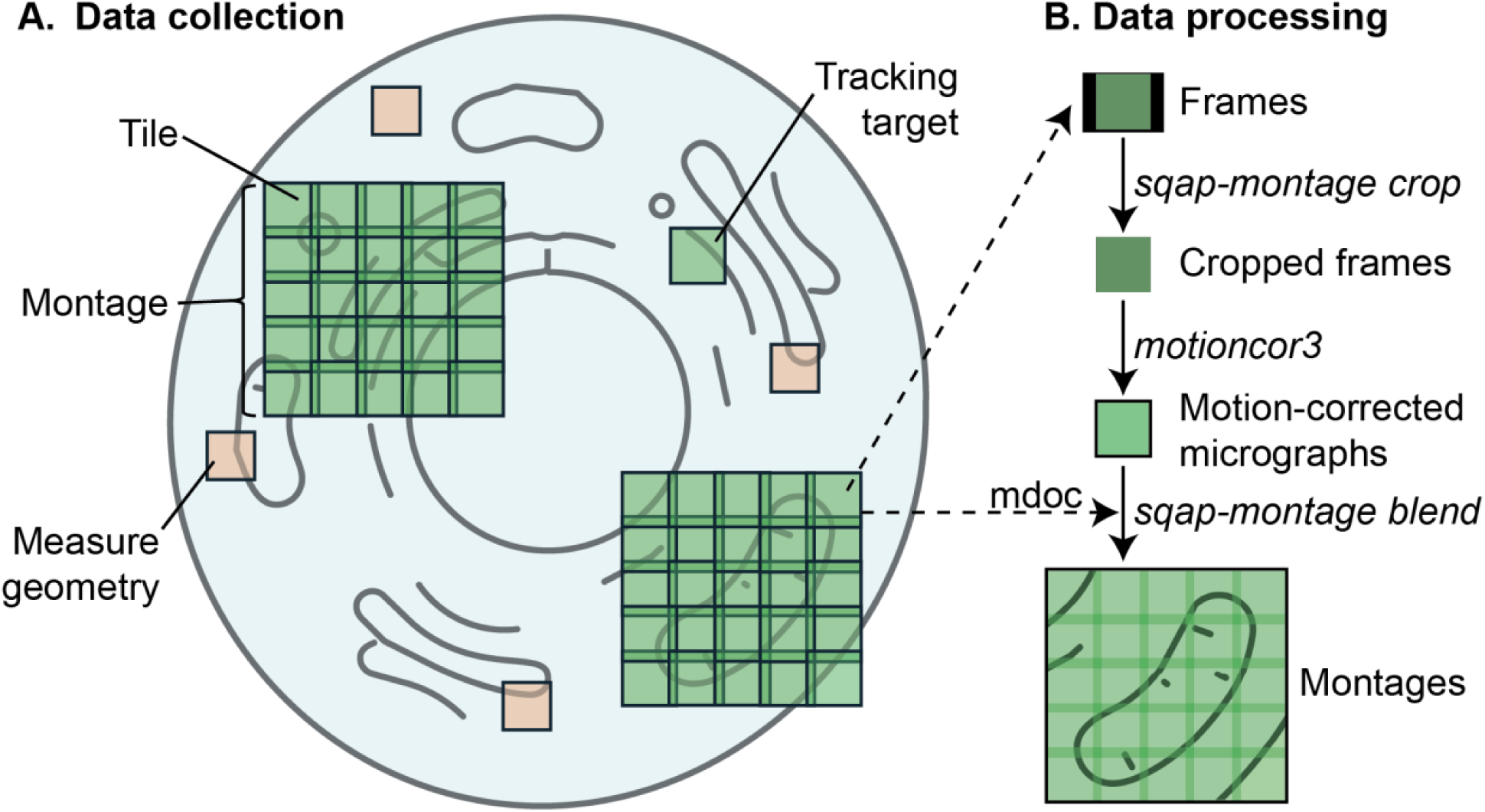
Workffow for square montage data collection and processing. (A) An example data collection strategy with PACE-tomo, showing the collection of two 5×5 square montages on a cell. A tracking target is established, and the orange targets are for geometry estimation. At each tilt, tiles are collected at each target location with a defined overlap to cover the regions of interest. We used this strategy to collect glial cell and malaria parasite data. The data collection and processing strategy for VLPs in holey grids is shown in Supplementary Figure 3. (B) An example data processing strategy for square beam montages. First, we cropped the frames for each tile. Then, the cropped frames were motion-corrected using MotionCor3. The motion-corrected and dose-weighted micrographs are then assembled into montages with the metadata in the SerialEM mdoc files.

We developed a Python pipeline, *sqap-montage*, to pre-process tiles into montage tilt series (Figure 1B). The pipeline has 3 core functions: *write-config*, *crop*, and *blend*. The *write-config* function creates a yaml file to define the directories and parameters for frame cropping and montage assembly. The *crop* function crops images to the illumination area. Users can detect the edges of the square beam using Otsu thresholding (Otsu, 1979) for automatic detection, or provide a user-defined threshold. Cropping can be done either before or after motion correction; it is important to run motion correction with patches (e.g. 3×3 or 5×5) so that the illuminated areas of the sensor are well corrected. The *blend* function calls the IMOD *blendmont* function (Mastronarde and Held, 2017) to align overlap regions and blend the tiles via interpolation. The function also writes out the tilt series stack, mdoc, and rawtlt files needed for downstream tilt series alignment in AreTomo (Zheng et al., 2022), RELION (Burt et al., 2024), or Warp (Tegunov and Cramer, 2019) (Figure 1B). Furthermore, the code is parallelized such that using 41 CPUs (1 per tilt image), montage frames and averages can be created in ∼15 minutes per tilt series.

### Improvements in square montages with overlapped data collection

The ideal data collection strategy for square beam montages is to collect tiles without any overlaps. However, in practice, we observed tiling inaccuracies that resulted in artifacts such as gaps between tiles, double imaging of the sample, and discontinuous features (Supplementary Figure 1A & B). These tiling inaccuracies could have arisen from sample drift during imaging or errors in the microscope’s beam-image shift. We found that we could produce montages without most of these artifacts by collecting data with 5–15% tile overlap, then aligning and blending the overlapping regions using IMOD’s blendmont (Supplementary Figure 1C–E).

We observed that alignment of tiles by blendmont was not perfect, especially at higher tilts where the signal-to-noise ratio decreases. We used IMOD’s Midas to examine the tile alignments and manually adjusted any poorly aligned tiles to produce a spatially coherent montage. While this step was important for good tilt series alignment of the montage, doing so is labor-intensive and time-consuming. Future work in this area could involve the use of contrast enhancement tools such as denoising (Bepler et al., 2020) to improve the signal for cross-correlation in the small overlap regions between tiles, or developing machine learning-based tools that can learn and improve alignments between tiles.

A data collection strategy with overlaps between tiles introduces areas on the montage with a higher accumulated dose. We simulated the dose distribution for square beams in a 3×3 with 10% overlap (Figure 2A), and for round beams with 10% overlap and a spiral translation data collection pattern as described in (Yang et al., 2023) (Figure 2B). In an ideal scenario (square montages with 0% overlap), the dose over the entire reconstructed area is evenly distributed at 123 e/Å^2^. Collecting square beam montages with a 10% overlap increases the average dose to 144 e/Å^2^, as compared to 211 e/Å^2^ for round beam montages with 10% overlap (Figure 2C). Although the maximum dose on square beam montages is higher than with round beam montages, the highly dosed areas are concentrated in more limited regions of the sample. This allows more of the sample to contain more acceptable doses: 53% of the sample is dosed at 123 e/Å^2^ for square beams with 10% overlap, compared to only 6% for round beams with 10% overlap (Figure 2C).

**Figure 2.**
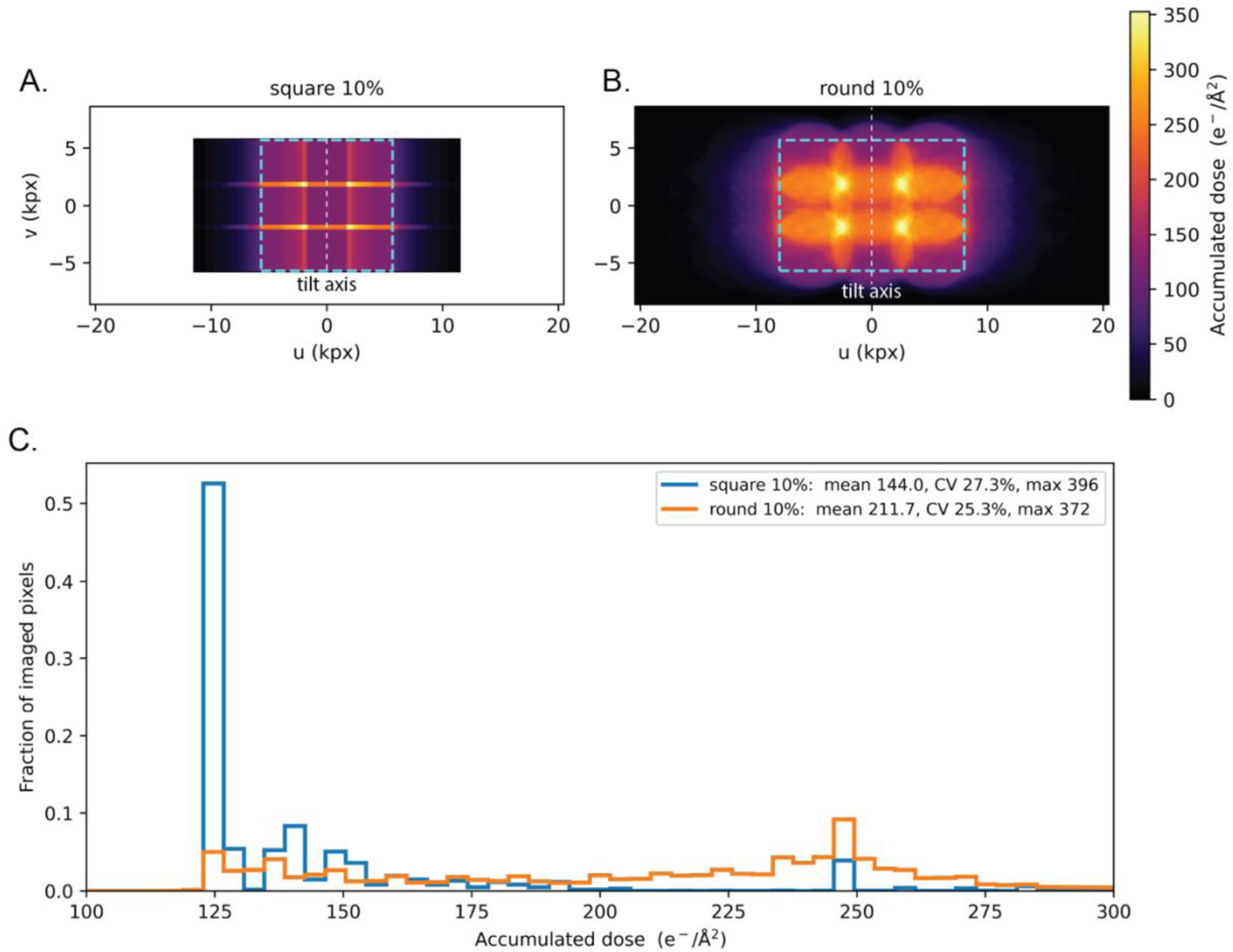
Dose distribution plots for montage tomography data collection schemes, comparing: (A) 3×3 square beam montage with 10% overlap between tiles, with (B) 3×3 round beam montage with 10% overlap between tiles, collected with a spiral pattern in the same manner described in Yang et al., 2023. Dotted squares indicate the total montage region captured by the sensor at zero tilt, which we also use to calculate the dose distribution histogram in C. Tilt axis in both A and B runs perpendicular to the u axis, through 0. The total exposed area for square beam montages is smaller than for round beam montages, shown by the smaller field of view of black pixels in A vs B. (C) Dose distribution histogram plot for square beam montage with 10% overlap (blue), and round beam montage with 10% overlap (orange). In an ideal scenario of square beam montages with no overlap, all pixels will have 123 e/Å^2^ accumulated dose. Legend: Mean = average electron dose per pixel; CV = coefficient of variation, the standard deviation divided by the mean; max = maximum electron dose received by a pixel.

We used Leopard-EM (Giammar et al., 2026) to perform high-resolution 2D template matching of apoferritin and found that regions in tiles previously imaged no longer contain high-resolution information (Supplementary Figure 2).

### High-resolution STA reconstruction obtained from square montages

To determine if square beam montages retain high-resolution information, we collected eight 3×3 montage tomograms of PP7 VLPs from a single square on a holey carbon grid (Supplementary Figure 3). Each montage covered a field of view of ∼1.2 by 1.2 µm, or ∼1.4 µm^2^ (Figure 3). The total time to collect eight 3×3 montages from one square and stage position was 3 hours and 5 minutes. From these montages, STA of 2,295 VLPs with icosahedral symmetry yielded a 3.5 Å reconstruction showing side-chain densities (Figure 3, Supplementary Figure 4).

**Figure 3.**
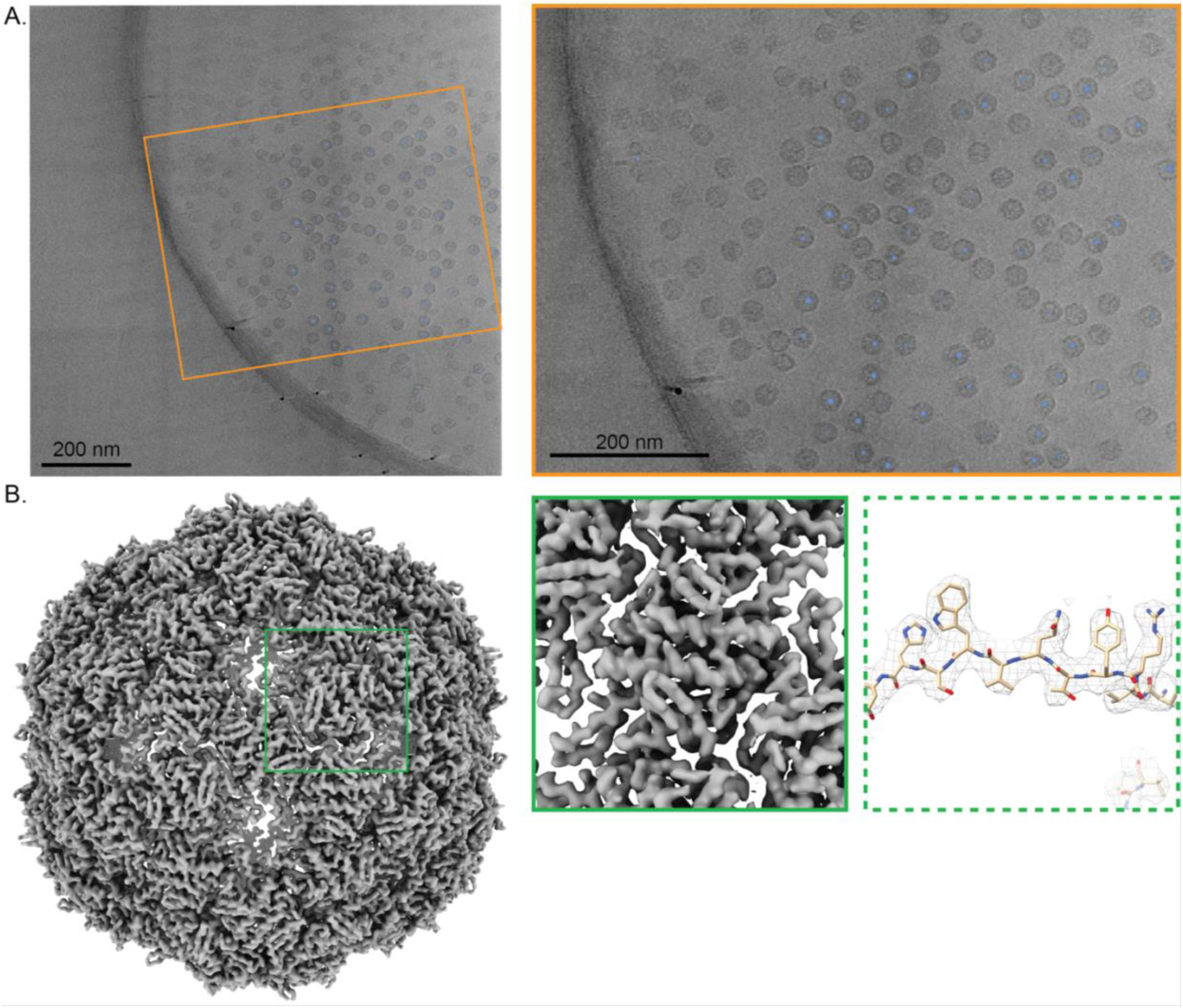
(A) A slice through a montage tomogram of PP7 VLPs on a holey carbon grid. Orange inset shows a zoomed-in region of the volume, with cryolo picks of VLPs shown as blue dots. (B) STA reconstruction of PP7 VLPs to 3.5 Å. A zoomed-in region of the VLP is shown (solid green box), and a β sheet with a model refined into the map (dashed green box). Fourier shell correlation (FSC) curve shown below.

We used our dose distribution simulator to partition VLP particles based on their location within the tomograms given our VLP data acquisition conditions. We separated the VLPs into two groups: one that received 123 e/Å^2^ total dose, and a second group that received more dose (>123 e/Å^2^), averaging around 150 e/Å^2^ (Supplementary Figure 5). The first group had 981 particles (∼43% of total particles) and reconstructed to 3.63 Å. The second group had 1,167 particles, of which 981 particles reconstructed to 3.85 Å. The slightly lower-resolution structure from particles in the higher-dose group is consistent with radiation damage-dependent changes to protein structural integrity.

We found that collecting slightly overlapping images is essential for producing a spatially coherent montage and tilt series, which in turn yields high-quality tomograms. Unlike round beams, square beams confine the high dose from overlaps to limited regions of the montage, so most of the field-of-view retains information conducive to high-resolution structure determination.

### Imaging large fields-of-view in lamellae with square montage tomography

Next, we applied square montage tomography to two distinct cellular samples to demonstrate its applicability for investigating biological systems of interest.

We collected a 5×5 montage of a glial cell in a neuronal cell culture grown on a grid (Figure 4). At a pixel size of 2.7 Å, the 5×5 montage covers an area of ∼5 x 5 µm. Our goal with imaging this cell was to characterize microtubule organization, which often extends several microns, well beyond the field-of-view of a single micrograph. To that end, we identified and segmented 52 microtubules across the entire montage. Of the 52 microtubules identified, the vast majority spanned the entire field of view, with only two microtubule ends visible. Measurements of the microtubules showed that 46 out of 54 microtubules (85%) exceeded 3 µm in length in this cell (Figure 4C). This contrasts with the shorter 2 µm average length observed in mammalian mitotic spindles (Kiewisz et al., 2022), but is consistent with reports of microtubules longer than 12 µm in mouse neuronal axons (Yu and Baas, 1994). We then used STA of 41,091 microtubule particles to determine their polarity and found ∼70% of the microtubule plus-ends point in the same direction. We envision that future work with square montages can include studying microtubule motor locations and their involvement in microtubule organization over long distances in cells (Conway et al., 2026).

**Figure 4.**
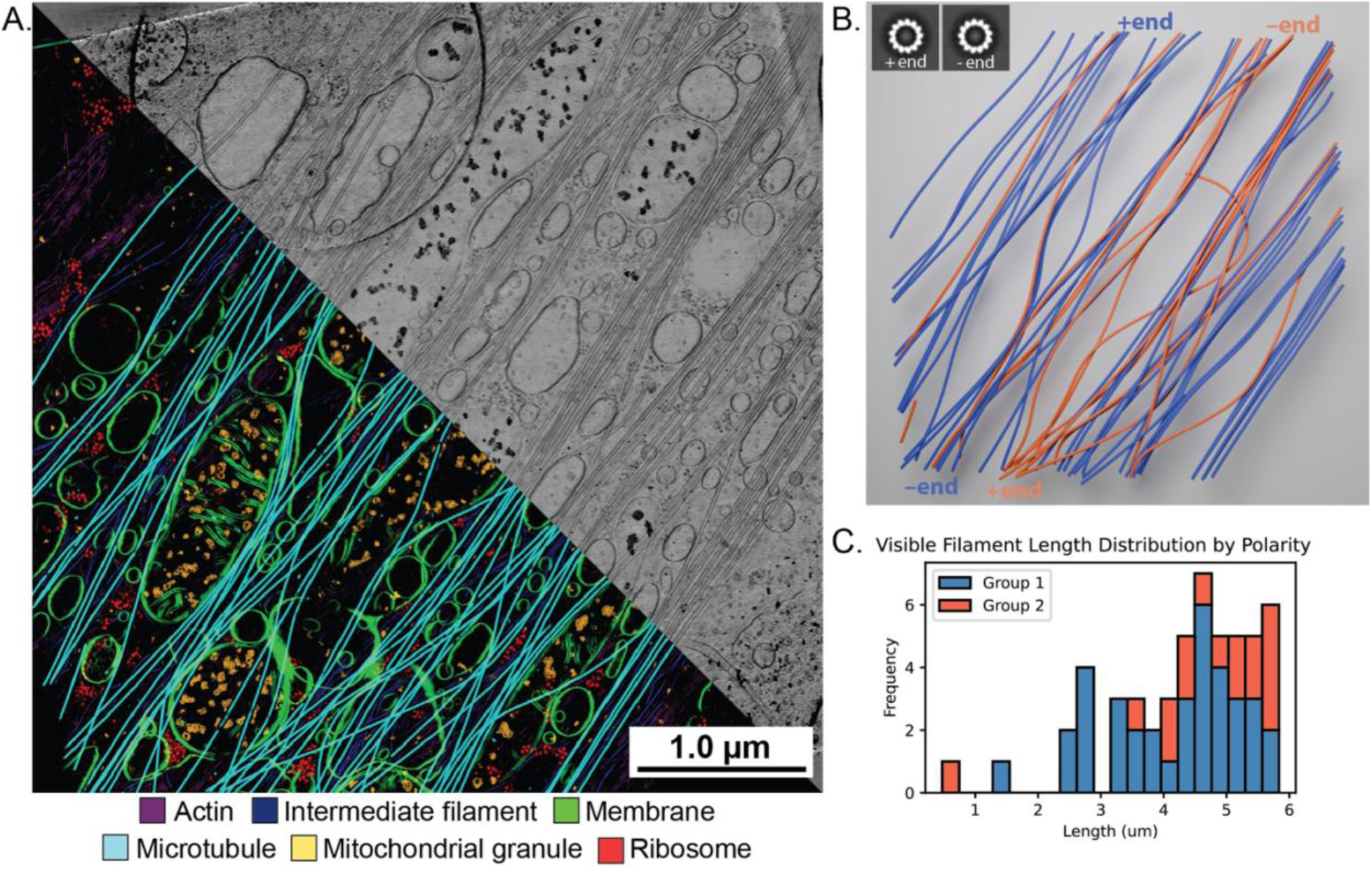
Segmentation and microtubule analysis of a 5×5 square montage of a glial cell. (A) Segmentations (bottom left) and a projection (top right) of the glial cell, covering an area of ∼5 x 5 µm. Cellular features were segmented with easymode and could be traced continuously across the montage. (B) Microtubules segmented with Tardis, colored by polarity, which was determined by sub-tomogram averaging. (C) Analysis of microtubule filament lengths visible in the montage.

We next imaged lamellae of schizont stage malaria parasites within their host red blood cells (Figure 5). During the schizont stage, a single parasite asexually reproduces into several daughter merozoites, which are then released to invade other red blood cells, thereby propagating infection and the symptoms of malaria. We collected a 7×7 montage covering an area of 5.1 x 5.1 µm (Figure 5A). Developing daughter merozoites are arranged around a central digestive vacuole that contains hemozoin crystals. Within each merozoite, organelles and other cellular structures are captured in context with one another while preserving molecular features, such as ribosomes and nuclear condensates (Figure 5B). Capturing intact schizonts in their entirety reveals the extensive cellular remodeling that underpins this critical stage of the malaria parasite lifecycle while retaining sufficient information for downstream high-resolution STA averaging of newly identified features.

**Figure 5.**
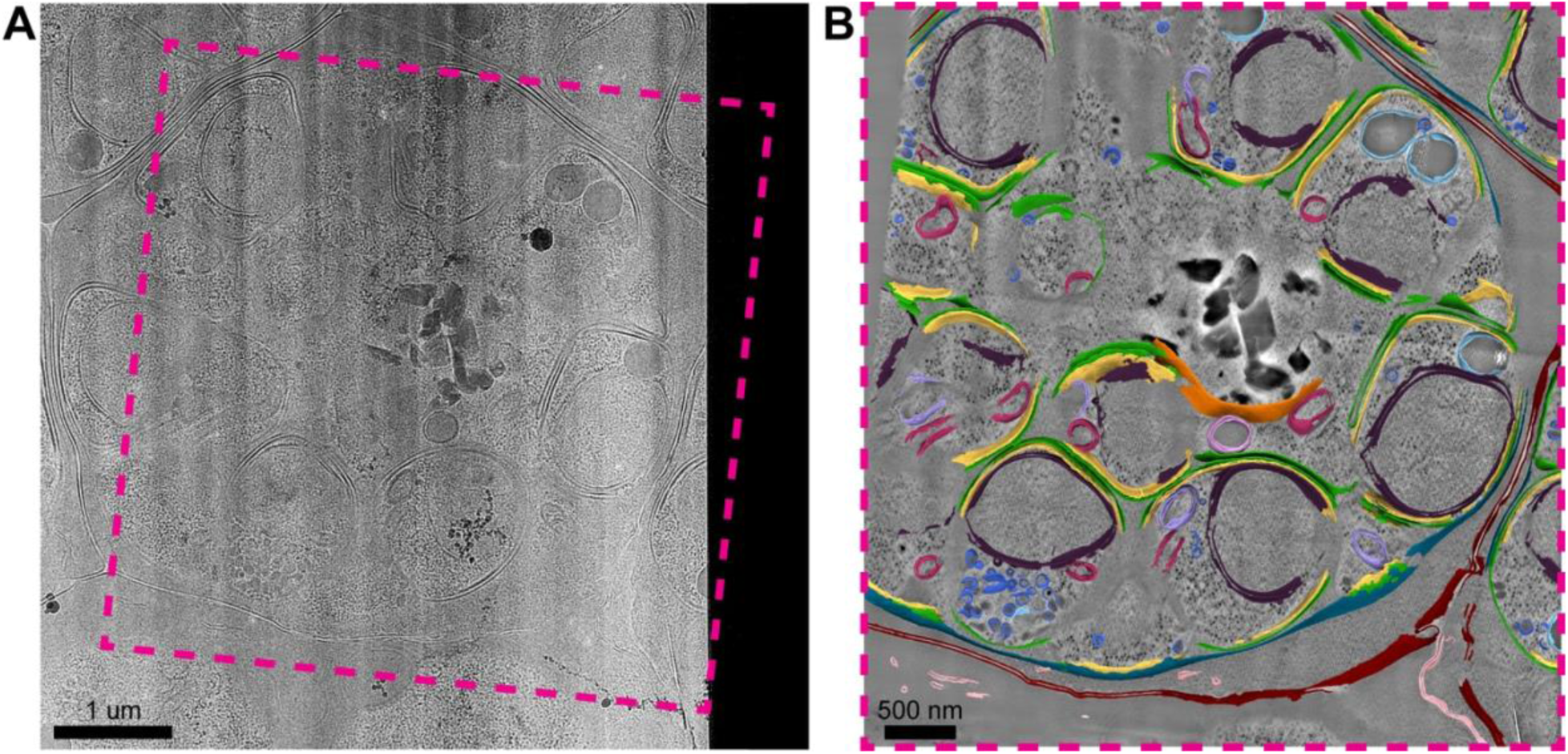
Square beam montage tomography of FIB-milled malaria parasites. (A) Low magnification overview image of lamella. Imaged region is indicated by magenta box. (B) High magnification 7×7 montage tomogram slice of region indicated in A. Membrane segmentations of erythrocyte membrane (red), parasitophorous vacuole membrane (teal), parasite membrane (green), inner membrane complex (gold), nuclei (purple), mitochondria (maroon), apicoplasts (lilac), rhoptries (sky blue), micronemes (blue), digestive vacuole (orange), hemoglobin vesicle (pink), and Maurer’s clefts (carnation).

Compared with workflows that use low-magnification overviews to capture contextual information (Gebauer et al., 2026; Watson et al., 2026), square beam montages offer two key advantages. First, some biological processes may not be visible at low magnification, so using low-magnification overviews to target regions of interest for high-magnification imaging may cause researchers to miss features that are not readily apparent. Second, collecting with square beams will allow researchers to gather data comprehensively then mine it for biological features of interest. These features may be located close to each other, such that researchers using round beams may have to choose between them to image at high magnification due to overlapping exposure areas and accumulating radiation damage; square beams will allow them to capture these features within a montage. Furthermore, the production of biological lamellae is time-consuming, labor-intensive, and expensive, and may involve rare samples or events, so capturing more of the sample can help maximize the information gathered from each lamella.

One major challenge we have encountered is the significant space and computational requirements for processing montage data. For example, an unbinned 7×7 montage tilt series with 41 tilts encompasses ∼120 GB of memory. After binning by a factor of 5 to ∼10 Å/pixel in float16 format, the resulting tomogram still occupies 11 GB, which is further propagated by downstream processing files. We ran into memory issues when trying to perform denoising or missing wedge correction on these montage tomograms, despite using GPUs with 48 GB of VRAM. This required dividing the montage tomogram into subvolumes that existing data processing tools could handle, then reassembling the resulting subvolume results into a montage. Future developments will include better processing strategies for reconstructing and managing large montage tomograms, and improved file and volume compression practices to reduce and optimize data storage usage (Fluty and Ludtke, 2022; Larsson, 2026).

## Conclusion

In this work, we demonstrate that square beam montage tomography provides both high-resolution information about a biological system and the large field-of-view for situating the biological context. Our workflow uses data processing tools already available to the cryo-ET community, enabling easy adoption and integration into existing workflows. Overall, we envision that the results and protocols presented here will enable scientists to implement and use square electron beams to image and study mesoscale processes at high resolution.

## Methods

### Sample and grid preparation

Grids for PP7 virus-like particles (VLP) (Keshavarz-Joud et al., 2025) and apoferritin were prepared as follows: Quantifoil R2/2 300 mesh Au grids were hydrophilized by plasma cleaning with a mixture of H2 and O2 gas (6.4:27.5 ratio) at 50 W for 30 s in a Solarus Model 950 Advanced Plasma System (Gatan). 3 µl of 8 mg/ml mouse apoferritin or 1 mg/ml PP7 VLP was pipetted onto the grid, blotted for 3-4 s in a Vitrobot at 20°C and 100% relative humidity, then vitrified in liquid ethane cooled to liquid nitrogen temperature.

Lamellae of malaria parasite were generated as previously described (Anton et al., 2025). In brief, asexual Exp2-mNG P. falciparum parasites were cultured in erythrocytes, and the schizont stage was enriched using a cushion of 65% Percoll and synchronized by treatment with 10 nM C1 (BEI Resources MRT-0207065) for 4 hours. Synchronized parasites at the schizont stage were pelleted at room temperature and resuspended in complete RPMI buffer with 10 nM C1. 3.5 μL of infected erythrocytes were applied to glow-discharged Quantifoil R2/2 Cu 200 mesh grids, back-blotted with Whatman Grade 1 filter paper, and plunge frozen in liquid nitrogen-cooled liquid ethane using a manual plunger. After clipping, grids were assessed for ice thickness and cell density using a Leica EM Cryo CLEM. Lamellae were milled on an Aquilos cryo-FIB/SEM (Thermo Fisher Scientific). Grids were sputtered with platinum metal and coated with an organometallic platinum layer using the gas injection system (GIS). Relief trenches were milled at 40° and subsequent lamella thinning was performed at 7°. Target lamellae sites were manually milled in parallel at 30 kV with steps of 0.5 nA, 0.3 nA, 0.1 nA, 50 pA, and 10 pA. Lamellae for cryo-ET were polished to a target thickness of 150 nm.

For the glial cell sample, E18 C57 dissociated mouse hippocampal neurons were obtained from Transnetx (SKU: C57EDHT). Cells were warmed up in the shipping media at 37°C for 10 minutes. Then, cells were diluted 6-fold in temperature- and gas-equilibrated (37°C, 5% CO2) plating media of Neurobasal further comprising 1X B27 (Gibco, A3582801), 50 μM Glutamax (ThermoFisher Scientific, 35050079), 25 μM glutamate, and 1X Penicillin - Streptomycin (Gibco, 15-140-148), further activated at 37°C for 10 minutes, then seeded onto plates. The plates were prepared by coating with 600 µL of 50 µg/mL poly-D-lysine (PDL) (Gibco, A3890401) overnight at room temperature, then coating with laminin (Gibco, 23-017-015) for 2 hours at 37°C right before seeding the neurons. During the PDL and laminin coating steps, 200 mesh R6/100 Quantifoil grids (Catalog number Q210CR6100) were placed inside the plates after plasma cleaning to coat the grids. Neurons were seeded directly on top of the grids, then kept in the incubator (37°C, 5% CO2) for 2 hours prior to being supplemented with additional plating media. Three days after plating, 750 mL of media was aspirated and replaced with culture media, which comprised the same components as the plating media except for glutamate. The media replacement was performed every 3 to 4 days for 3 weeks until freezing. Grids were frozen by plunge-freezing into liquid ethane on a Leica GP2 plunge freezer after 2 seconds of back blotting.

### Data collection

A detailed protocol for hardware and software setup and data collection is available on protocols.io. Here, we summarize the process.

For STA, PP7 VLP grids were used. Data were collected with PACE-tomo at a pixel size of 1.038 Å/px, exposure time of 0.24 s, for a total dose of 3.26 e/Å^2^ per tilt distributed over 4 frames, a dose-symmetric tilt range of −60° to 60°, in 3° increments, for a total of 41 tilts and a total dose of 134 e/Å2 per tilt series. At each stage position, the central hole with VLPs served as the tracking target, and 3×3 montages were collected from the surrounding eight holes, with an 8% overlap between tiles.

For 2DTM, apoferritin grids were used. Data were collected on a Cs-corrected microscope by using PACE-tomo at a pixel size of 1.076 Å/px, exposure time of 4.94 s, for a total dose of 46.54 e/Å^2^ distributed over 33 frames at 0° tilt. A 3×3 montage was collected at each target location with a 20% overlap between tiles.

For segmentation, a glial cell from a neuronal cell culture and malaria parasite lamellae were used. For the glial cell, data were collected on a Cs-corrected microscope with PACE-tomo at a pixel size of 2.709 Å/px, exposure time of 0.6 s, for a dose of 3.1 e/Å^2^ per tilt distributed over 12 frames, a dose-symmetric tilt range of -60° to 60°, in 3° increments, for a total of 41 tilts and a total dose of 127 e/Å^2^ for the tilt series. A 5×5 montage was collected with a 10% overlap between tiles. For the malaria lamellae, data were collected on a Cs-corrected microscope with PACE-tomo at a pixel size of 1.91 Å/px, exposure time of 0.33 s, for a dose of 3.02 e/Å^2^ per tilt distributed over 10 frames, a dose-symmetric tilt range of -54° to 66°, in 3° increments, for a total of 41 tilts and a total dose of 123 e/Å^2^ per tilt series. Data collection started at 6 deg to account for lamella pre-tilt. 7×7 and 11×11 montages were collected on these lamellae with 8% overlap between tiles.

A summary of data acquisition parameters can be found in Supplementary Table 1.

### Data pre-processing

Frames were first cropped to remove the unilluminated edges. Motion correction was then performed on the cropped micrographs with motioncor3 (Zheng et al., 2017). VLP montage tilt series were assembled with IMOD’s blendmont function to align overlap regions and blend them via interpolation (Mastronarde and Held, 2017). For malaria lamellae data, tiles from each tilt were additionally first normalized by histogram matching to the central tile before blending. Doing so improved the contrast in the resulting tomograms.

### Sub-tomogram averaging

STA of VLPs was performed in RELION-5.0 (Burt et al., 2024). VLP montage tilt series were first padded so that all montage tilts had the same dimensions. These tilt series were imported into RELION-5.0 as motion-corrected micrographs. mdoc files were updated to change each ZValue’s SubFramePath to the appropriate montage tilt micrograph name and used as input for RELION-5. CTF estimation was performed, and montage tilts with bad CTF estimates were manually removed. Remaining tilts were aligned with RELION’s wrapper for AreTomo2 (Zheng et al., 2022) using an estimated thickness of 200 nm. Tomograms were reconstructed with Xdim = 14000, Ydim = 13000, Zdim = 2000, and binned to 10 Å/px.

A sampling of particles was manually picked from the tomograms and then used to train cryolo (Wagner et al., 2019) to perform comprehensive picking across all tomograms. Cryolo’s output particles were then converted to RELION’s coordinate system with a script adapted from C.F. David Hou (https://forum.image.sc/t/convert-cbox-3d-cbox-to-relion-tomo-particles-star/95797), then re-imported into RELION. Particles were extracted as 2D stacks, then after a 3D classification and refinement job, four rounds of iterative CTF refinement, Bayesian polishing, and 3D refinement were done to obtain the final reconstruction at 3.5 Å. The crystal structure of PP7 VLP (1DWN) was fit into the final map and refined in real space with Phenix (Liebschner et al., 2019).

The same data were processed with WarpTools (Tegunov and Cramer, 2019) to yield a 3.5 Å reconstruction.

### Electron dose distribution simulation

An electron dose distribution simulator was designed with Claude. A thin specimen is assumed. The electron beam travels down the optical axis, z, and encounters a specimen with the frame (u, v), where u is the axis perpendicular to the tilt axis and v the axis parallel to the tilt axis. A tilt range of -60° to 60° is defined, with 3° steps, for a total of 41 tilts, and a dose of 3 e/Å^2^ per montage tile per tilt. A square beam was simulated at the size of the short dimension of a K3 detector (4092 pixels), and a 3×3 montage acquisition pattern with 10% overlap between tiles was set. The same beam-image shift vectors were maintained throughout the tilt series such that the beam positions are fixed relative to each other while the sample was being tilted, in a manner identical to our data collection strategy with PACE-tomo. The dose on the specimen was plotted as a heat map. Round beam simulations were performed as described in (Yang et al., 2023). Briefly, round beams with a diameter exactly encompassing a K3 detector (5760×4092 pixels) were simulated, and a 3×3 montage acquisition pattern with 10% overlap between tiles was requested. At each tilt, the beam was translated in an Archimedean spiral pattern with a growing radius to evenly distribute the dose across the sample.

### Tomogram reconstruction and segmentation

For glial cell data, motion-corrected tiles were first normalized, then blended to form the montage. Errors in tiling during montage assembly were manually corrected using IMOD’s Midas. The montage tilt series was imported into WarpTools (Tegunov and Cramer, 2019). The tilt series was aligned with the WarpTools wrapper for etomo patch tracking (Mastronarde and Held, 2017). The tilt series alignments were then improved with MissAlignment (Chaillet et al., 2026). The CTF and defocus handedness were estimated, then the tomogram was reconstructed with WarpTools. tomosplit.py was used to divide the montage volume into sub-volumes. Each sub-volume was segmented with easymode (So-Last et al., 2026), and the resulting segmentations reassembled into the montage with tomosplit.py. Each sub-volume was also denoised and missing wedge corrected with IsoNet2 (Liu et al., 2025), and the results reassembled into the full denoised and missing wedge corrected montage with tomosplit.py. The resulting segmentations and volumes were visualized with napari (Sofroniew et al., 2026).

Microtubules in the glial cell data were additionally segmented with Tardis (Kiewisz et al., 2024). The segmentation result from Tardis was manually corrected with Dragonfly (Gendron et al., 2021). Using the resulting binary mask of microtubules, coordinates along individual microtubule filaments were extracted at 4 nm steps with Warp (Tegunov and Cramer, 2019) and aligned for sub-tomogram averaging using RELION (Burt et al., 2024). The polarity of the microtubules was assigned by first evaluating the clockwise versus counterclockwise direction of the slew of the averaged structure, then mapping the Euler angles of the particle back to the tomogram. For all microtubule filaments, >90% of the polarities assigned agreed, indicating high confidence in the overall polarity estimation.

For malaria parasite data, motion-corrected tiles were first normalized then blended to form the montage. Errors in tiling during montage assembly were manually corrected using IMOD’s Midas. The tracking tilt series was used to determine the tilt axis rotation value in etomo (Mastronarde and Held, 2017). AreTomo3 was then used to determine the defocus handedness and refine the tilt axis rotation value (Peck et al., 2025). Unmilled edges of lamella in the montage tilt series were replaced with background by ccderaser to improve tilt series alignment. Montage tilt series were imported into WarpTools (Tegunov and Cramer, 2019). Tilt series alignment was performed using AreTomo2 with local patches matching the montage tile pattern through the WarpTools wrapper and improved with MissAlignment (Chaillet et al., 2026). Tomograms were leveled and then reconstructed at 10 Å/px. Tomograms were divided into patches using trimvol prior to even-odd tilt series processing in IsoNet2 (Liu et al., 2025). Segmentations of the IsoNet2-processed tomogram patches were generated in MemBrain-Seg (Lamm et al., 2024). Montage tomograms and segmentations were re-assembled with assemblevol. Tomograms and segmentations were visualized in 3dmod (Mastronarde and Held, 2017), ChimeraX (Meng et al., 2023), Mosaic (Maurer et al., 2025), and ARTIAX (Ermel et al., 2022).

## Supporting information

Detailed Protocol

## Data availability

Raw movie frames of VLPs and apoferritin collected for STA and 2DTM have been deposited in EMPIAR under the accession code EMPIAR-XXXXX. Accompanying STA VLP reconstruction has been deposited in EMDB under the accession code EMD-XXXXX. Raw movie frames of the glial cell have been deposited in EMPIAR under the accession code EMPIAR-XXXXX.

## Code availability

PACE-tomo code is available at https://github.com/eisfabian/PACEtomo. Square montage data processing code is available at https://github.com/hamid13r/sqap-montage. Dose distribution simulator and tomosplit.py are available at https://github.com/eydchua/sqap-montage-accessories.

## Acknowledgements

We thank M. Kikkawa (University of Tokyo) for the apoferritin plasmid and B. Kloss (NYSBC) for the expression and purification of apoferritin. We gratefully acknowledge J. Pellman and the Research Computing team (NYSBC) for their exceptional support for computing software and infrastructure. We thank A. Fitzpatrick (Columbia University) for providing access to his TFS Aquilos cryo-FIB-SEM.

## Funding

This work was supported by the Simons Electron Microscopy Center and Simons Resource for Automated Molecular Microscopy located at the New York Structural Biology Center, supported by grants from the Simons Foundation (SF349247), the NIH National Institute of General Medical Sciences (GM103310), and the NIH National Institute of Neurological Disorders and Stroke (NS125674-02).

## CRediT author statement

EYDC: conceptualization, methodology, software, validation, formal analysis, investigation, visualization, writing – original draft, writing – review & editing, project administration

HR: methodology, software, investigation, validation, writing – review C editing

JZ: methodology, visualization, investigation, writing – review C editing

FE: methodology, software, writing – review C editing

JDJ: investigation, software

YHS: investigation, resources

HW: resources

LMA: methodology MK: resources

CMH: supervision, funding acquisition, writing – review C editing

DG: supervision, funding acquisition, writing – review C editing

AdM: supervision, funding acquisition, writing – review & editing, project administration

## Notes

### Competing Interest Statement

EYDC and LMA are listed as inventors on a patent on using square beams in TEMs, publication number 20250166956.

