## Supplemental Figures and Table for "Imaging large fields-of-view at high resolution in cryo-ET with square beam montaging"

### Supplementary material

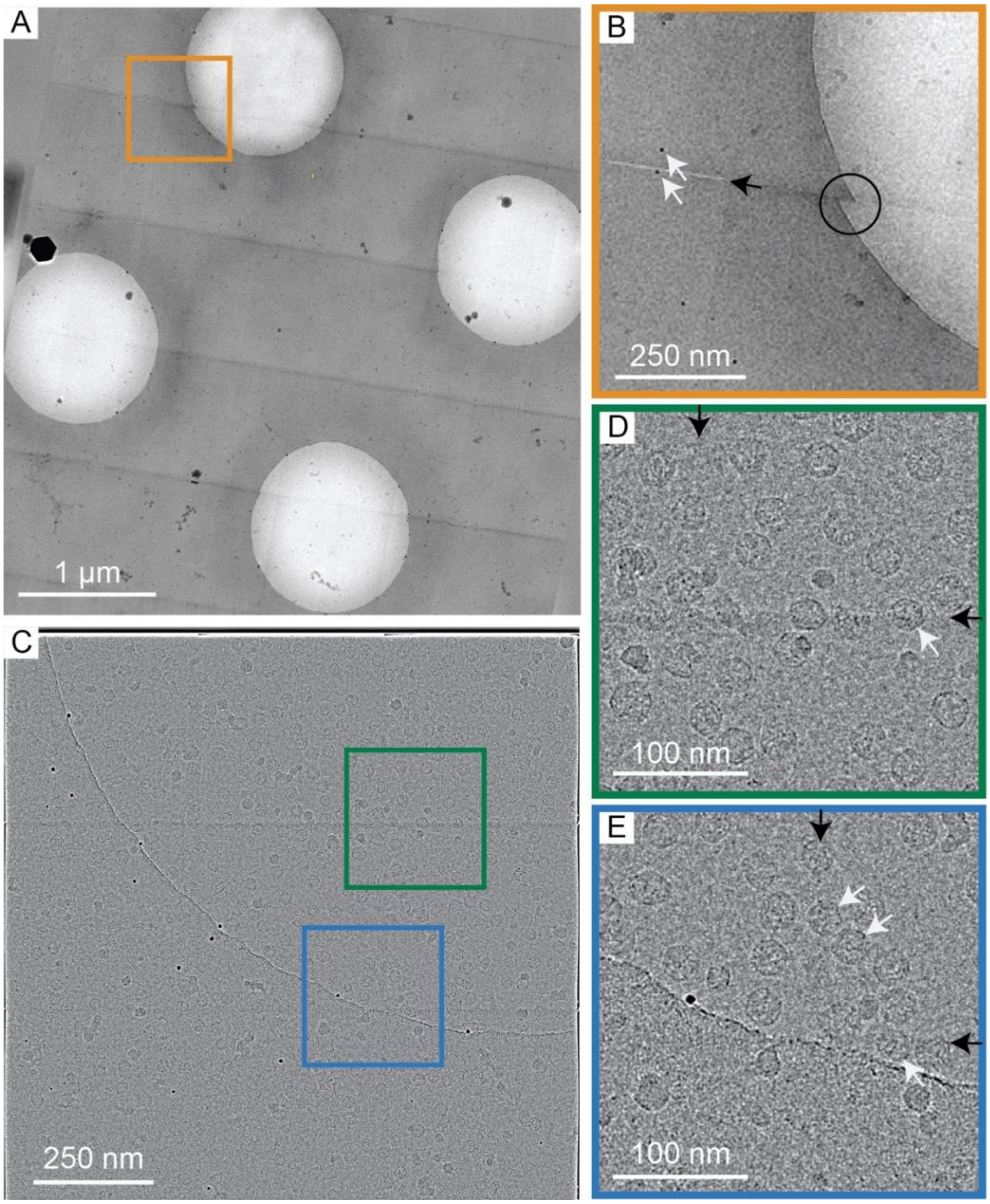


**Supplementary Figure 1.** (A) 5x5 montage of apoferritin on a holey carbon grid, collected with an end-to-end tiling strategy with no overlaps between tiles. The montage was assembled by butt-joining tiles. No alignment was performed prior to montage assembly. Image adapted from Chua et al (2024). (B) Zoomed-in region of A, showing a feature that was imaged in both tiles (white arrows), a gap between tiles (black arrow), and a discontinuous feature between tiles (black circle). (C) A 3x3 montage of VLPs on a holey carbon grid, collected at a higher magnification, with an 8% overlap between tiles. Tiles were aligned and blended with IMOD's blendmont function. (D) and (E) Zoomed-in regions of C at the blended regions showing visually-whole VLPs located in the overlap regions (white arrows), and a visible stitching line (black arrow).

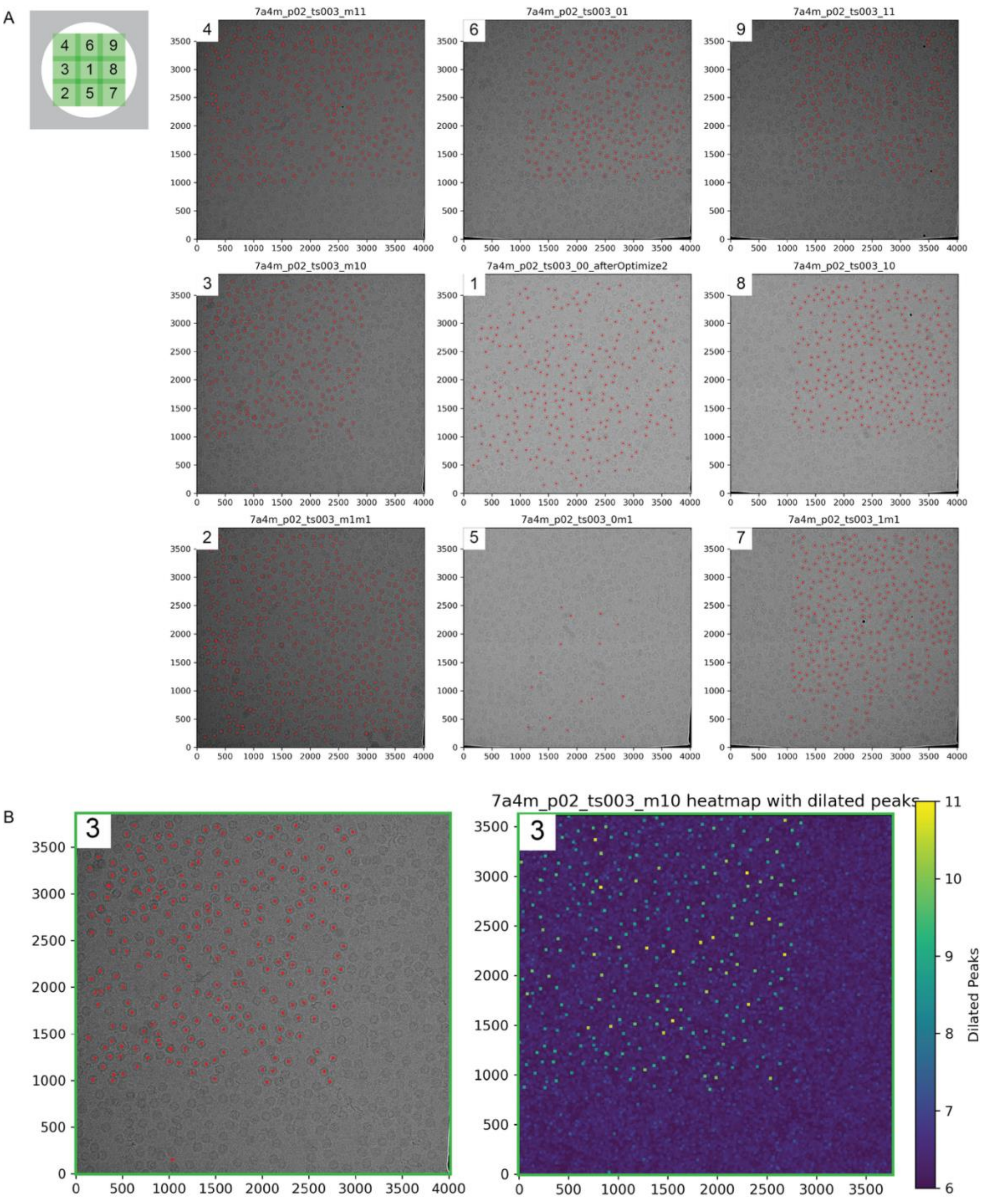


**Supplementary Figure 2.** Apoferritin particles identified by high-resolution 2D template matching with Leopard-EM within tiles collected for a 3x3 montage. (A) Tiles were collected with a 20% overlap. The schematic in the top left shows the data collection strategy, with numbers corresponding to the order in which the micrographs were collected. Each corresponding micrograph is also so labeled. (B) An enlarged version of tile #3, showing, on the left, 2DTM matches of apoferritin particles, and on the right, scaled maximum intensity projection (MIP) scores from the same micrographs, with the peaks dilated for visibility.

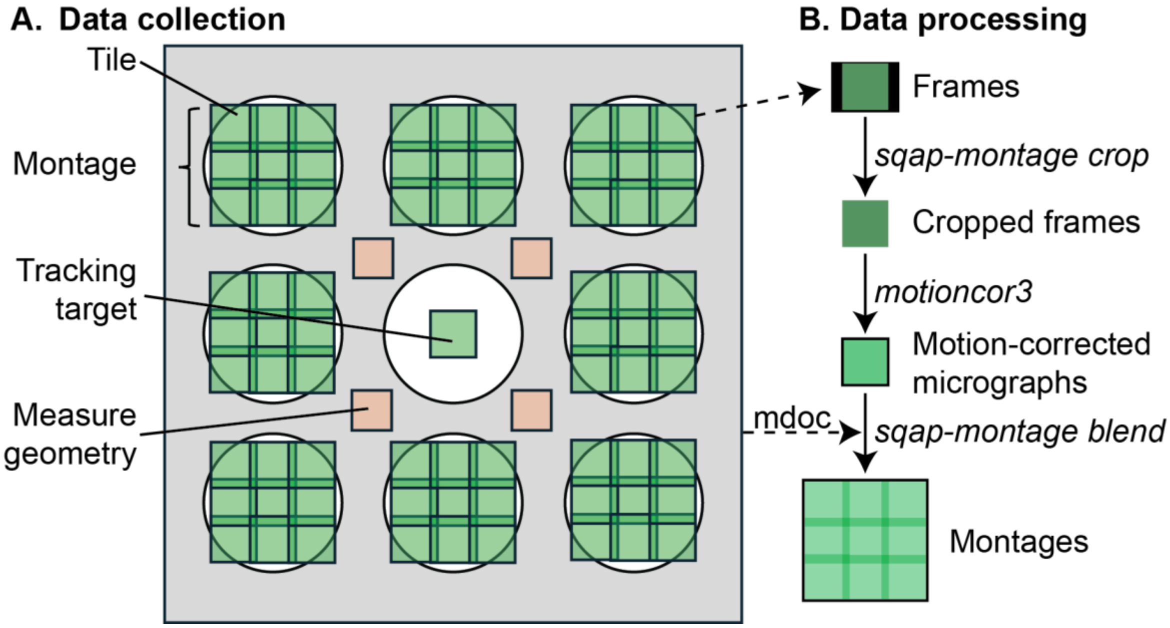


**Supplementary Figure 3.** (A) Square montage data collection strategy for PP7 VLPs on a holey carbon grid. The central hole was used as a tracking target, with geometry points set in orange. 3x3 montages were collected on the surrounding holes at each tilt, yielding 8 montages. (B) Data pre-processing pipeline for square montages. First, we cropped the frames to remove any unilluminated regions of the sensor, then the cropped frames were motion corrected with *motioncor3*. The motion corrected micrographs were then assembled into montages using the IMOD *blendmont* function together with metadata from PACE-tomo and the SerialEM output mdoc files.

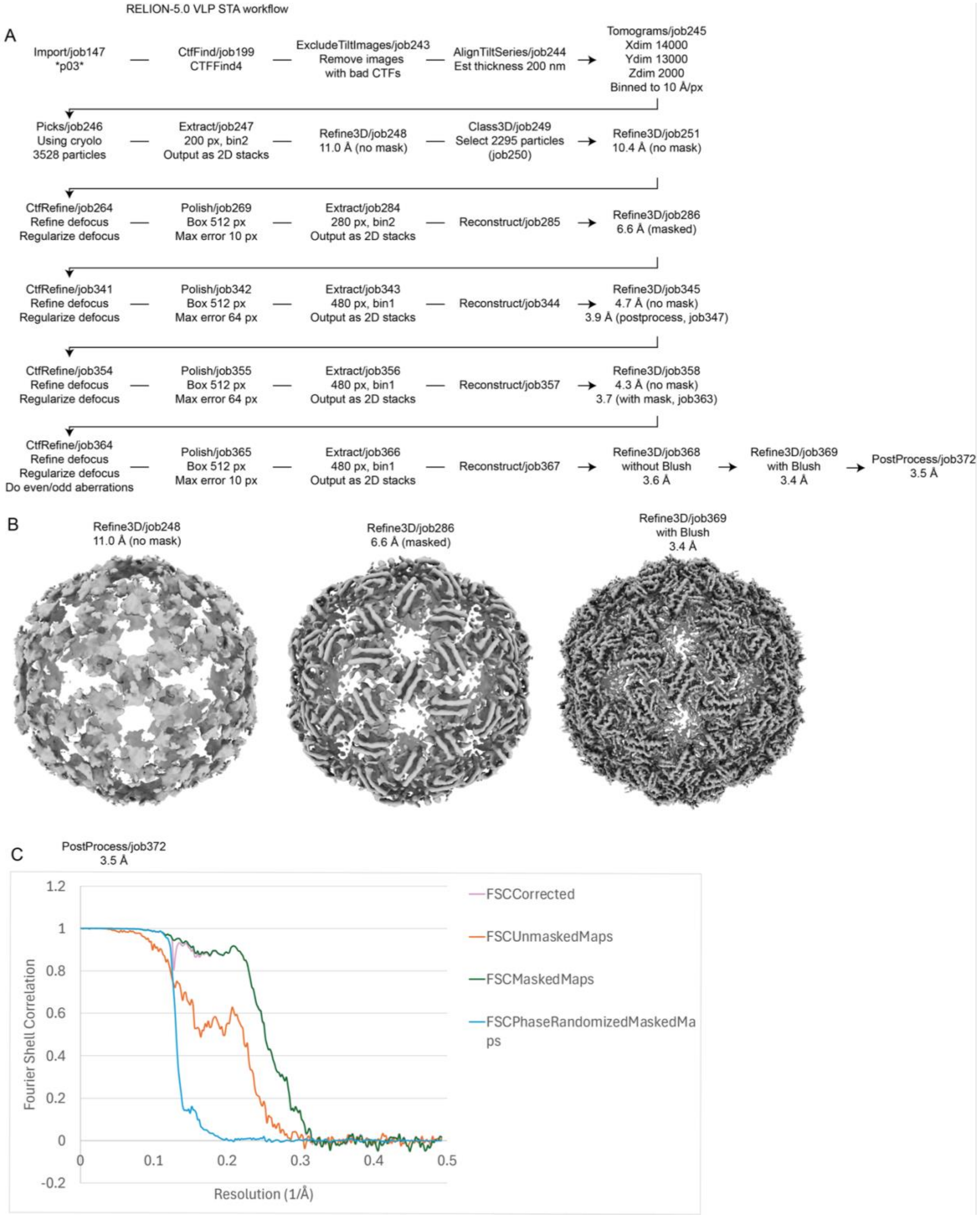


**Supplementary Figure 4.** (A) STA data processing workflow in RELION-5.0. Montage micrographs were imported into RELION then CTF estimated. Tilts with bad CTF estimates were manually excluded, then the remaining stack of tilts were aligned and reconstructed into tomogram volumes. A subset of particles were manually picked, which was then used as a training set for comprehensive picking in cryolo. The particles were extracted, refined, and classified in 3D for one round. Then, four rounds of CTF refinement and Bayesian polishing was done to refine the particles from 10.4 Å to a final of 3.5 Å. (B) Example maps from various stages of the STA workflow. (C) Fourier shell correlation (FSC) curve from the final postprocessing job.

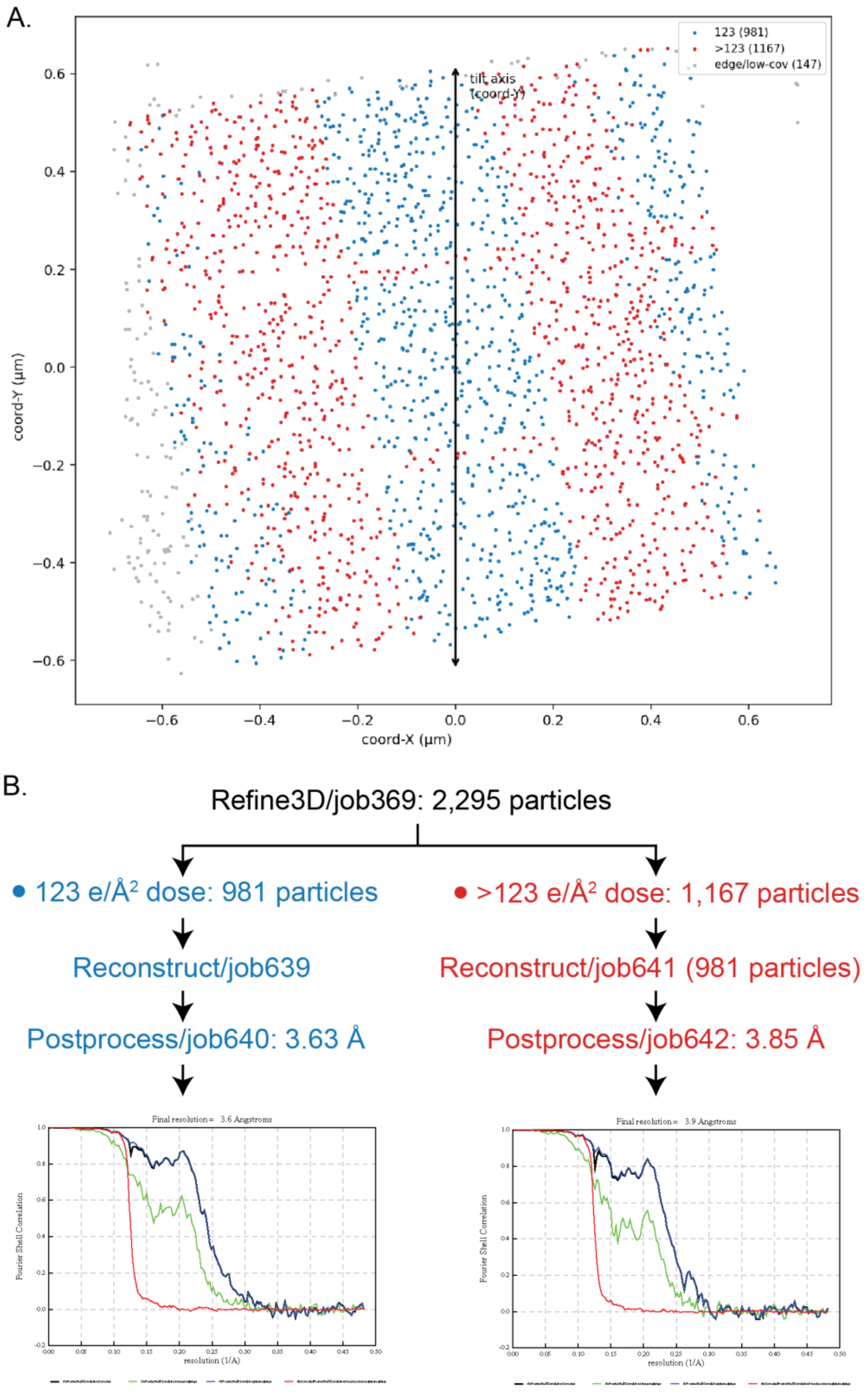


**Supplementary Figure 5.** (A) Plot of VLP locations within all eight 3x3 montage tomograms colored by whether they received 123 e/Å$^2$ total dose (blue), or more than 123 e/Å$^2$ total dose (red). Montage data collection conditions were simulated in the dose distribution simulator, then particles were partitioned according to their total dose received as determined by their location within the reconstructed tomogram. (B) Workflow for comparing particles receiving different amounts of dose. Particles in A were obtained from the final refinement job (see Supplementary Figure 1), then partitioned according to our simulations of electron dose across the samples.

| Application | STA | 2DTM | Segmentation | Segmentation |
|---|---|---|---|---|
| Sample | PP7 VLP | Mouse apoferritin | Glial cell | Malaria lamellae |
| **Microscope parameters** | | | | |
| Microscope | Titan Krios 1 | Titan Krios 2 (Cs-corrected) | Titan Krios 2 (Cs-corrected) | Titan Krios 2 (Cs-corrected) |
| Camera | Gatan K3 | Gatan K3 | Gatan K3 | Gatan K3 |
| Pixel size (Å/px) | 1.038 | 1.076 | 2.709 | 1.91 |
| **Tilting parameters** | | | | |
| Exposure time per tilt/image (s) | 0.24 | 4.94 | 0.6 | 0.33 |
| # frames per tilt/image | 4 | 33 | 12 | 10 |
| Dose per tilt/image (e/Å$^2$) | 3.26 | 46.54 | 3.1 | 3.02 |
| Tilt scheme | Dose-symmetric | No tilting | Dose-symmetric | Dose-symmetric |
| Tilt range | −60° to 60° | 0° | −60° to 60° | −54° to 66° |
| Number of tilts | 41 | 1 | 41 | 41 |
| Total dose (e/Å$^2$) | 134 | 46.54 | 127.1 | 123 |
| Montage size | 3x3 | 3x3 | 5x5 | 7x7 and 11x11 |

***Supplementary Table 1.*** *Summary of data collection parameters. Titan Krios 1 and 2 are both equipped with fringe-free illumination.*
