## Supplementary material for "Imaging large fields-of-view at high resolution in cryo-ET with square beam montaging": Detailed Protocol

### Protocol Supplement

The final version of this protocol will be uploaded on protocols.io.

### Microscope setup

#### A. Aperture preparation

The square apertures installed on our microscopes were purchased from Agar Scientific (product number AGAS3005P). A 50 µm square aperture was installed into the C2 aperture strip of a Titan Krios microscope (ThermoFisher Scientific). Before its installation into the microscope, the square aperture was plasma cleaned to remove any impurities, then maintained in a sealed container for several days to allow the charge to dissipate, allowing an easier insertion into the aperture strip.

On our Titan Krios microscope, we find that 50 µm is the largest aperture size that should be used. This is because the beam is condensed to the size of the sensor during imaging, which takes it to the bottom end of the microscope's parallel range at ~1 Å/pixel. With larger aperture sizes, the beam may no longer be parallel at the intensity where it is condensed to the size of the sensor.

| ℹ NOTE | Square or rectangular apertures can be used, depending on the configuration of your microscope sensor. Hamish Brown and team have developed apertures containing both square and rectangular holes at varying orientations that have the advantage of not requiring P2 lens alignment, which requires more intervention from a microscope engineer. |
|---|---|

#### B. Aperture change

| ⚠ WARNING | Changing the C2 aperture on a TEM should be done by a trained field service engineer following the protocol provided by the TEM-supplying company. Only a rough guide is provided here. |
|---|---|

Generally, the steps for C2 aperture change are as follows:

##### B-1. Prepare the Column for Aperture Change

1. Close column valves.
2. In the user interface (UI), navigate to Vacuum → Status and confirm that column vacuum ≤ $5 \times 10^{-5}$ Pa (typical operating vacuum ≈ $5–10 \times 10^{-7}$ Pa). Note the base pressure vacuum before starting the C2 aperture change in the service log.
3. Warm up the column by going to All Room temperature state.
4. Vent the octagon by pressing 'column vented' in the UI.

##### B-2. Exchange Aperture Strip

1. Remove the Xray protection cover on the C2 aperture strip, note the screws are spring loaded. ThermoFisher-supplied specialty screwdriver needed to remove the protected screws.
2. Unscrew the aperture rod lock nut/collar. Gently and smoothly withdraw the aperture rod in a straight, level motion. Note which side is up and down.
3. Place the removed aperture rod on a clean lint-free work surface.
4. Loosen the retaining screws on the aperture rod top with a Torx T6 screwdriver.
5. Using electrostatic-discharge-safe tweezers, remove the old aperture strip from the holder and place it in an appropriate container. Replace with the new aperture strip containing the square aperture or exchange one of the factory-supplied apertures in the existing aperture strip with a square one
6. Replace the screw and check that the rod tip and aperture strip are free from any visible debris or contamination.
7. Insert the aperture rod back into the microscope port and tighten the lock nut/collar.

##### B-3. Re-Establish Vacuum Conditions

1. In TUI, select 'all vacuum' to start pumping down the octagon and C2 aperture port.
2. Monitor the vacuum status and wait for full column vacuum to recover to $3x10^{-3}$ Pa.
3. Once the vacuum trip level is reached the system can be cooled down to LN2 state to improve the vacuum. Be sure to cryo-cycle overnight after the procedure is finished.

#### C. Square aperture alignment

| | |
|---|---|
| ⚠ **WARNING** | Changing the power in the P2 lens should only be done by a trained field service engineer following the protocol provided by the TEM-manufacturing company. Only a rough guide is provided here. |

##### C-1. Center and align C2 aperture

1. In the service application called 'Tools\AAS\Aperture alignment Wizard' rename the appropriate Aperture to, e.g. '50sq' to indicate both the width of the aperture and its square shape.
2. This application also allows centering of the apertures with regard to a condensed beam.

##### C-2. Align square beam to the sensor using the P2 lens

| | |
|---|---|
| ℹ **NOTE** | Changing the defocus, intensity, and P2 lens values will cause the square beam to rotate. This is because electrons spiral down the TEM column due |

| | |
|---|---|
| | to the effect of the lens's magnetic fields on moving charges, called the Lorentz force. |

1. Set the magnification that you will use for imaging, then condense the square beam such that it is approximately the size of the detector's short edge and set the spot size so that the beam gives a reasonable dose on the camera.
    a. There are two options for saving the magnification series for square beams: one is to set up new magnification values, and the other is to adapt an existing magnification value that is not commonly used.
    b. If setting up a new magnification series, then start with the equivalent round beam magnification.
    c. If adapting an existing magnification, then switch to that magnification to start.
2. If the beam does not fall square onto the camera, the P2 lens power needs to be changed to rotate the beam to fall square on the camera.
    a. In the UI, go to Setup → Free Lens Control. Record the baseline P2 value.
    b. If the UI does not have Free Lens Control, change the P2 lens with the optics control menu (see points f-k below otherwise skip steps f-k).
    c. Unlock the P2 value for manual adjustment. If you have a Gatan K3 camera on a separate computer, it may be helpful to open up a live view of the beam. If you have a Falcon camera, operate it in "continuous" mode with the Preview preset in SerialEM to do the P2 adjustment.
    d. Adjust the P2 value in one direction and check the direction of rotation of the square on the sensor. Adjust the P2 value in the direction that requires the least amount of change.
    e. Adjust the P2 value until the beam lands square onto the sensor. Iteratively adjust the intensity such that the square is the same size as the sensor, and the P2 values such that the beam lands square onto the sensor.
    f. Start the Windows Registry Editor and navigate to the key 'SMM'. Change it to '1'. This will show the micro map regardless which user is logged in (i.e. ShowMicroMap is true).
    g. In case the key is missing (as is on many Glacios systems) the key can be manually added.
    h. Restart windows and temserver to make the registry change active.
    i. Exit the Microscope software launcher (MSL) by right clicking and selecting exit.
    j. Start the MSL in customer service by right mouse click and select 'different user'
    k. In the software launcher a new menu is visible, Tools\Optics. Select menu [TEM optics] > P2 lens.
        i. Note that there is no 'Save' button in the micromap. Go to Direct Alignments > Beam Shift, adjust the P2 lens, adjust the beam shift,

then click ‘done’ in Direct Alignments to save the new P2 for that magnification.

l. In the microscope UI select the Mag that will be used for data collection
m. Set Gatan DM or Microscope companion for Falcon 4i to live view so the aperture is displayed live on the camera.
n. Change intensity to the dose needed. Note that the image rotates. It always does, but now this is visible with a square aperture.
o. Move the slider on the P2 dial in temspy till the C2 aperture square is aligned with the camera. The GUN lens might need to be changed if the needed dose cannot be set.
p. Hit save in the TEMspy dial box.

3. Repeat for each magnification you want to use the square aperture for.

### SerialEM setup

#### A. SerialEM

1. Download and install the latest version of SerialEM.
2. Install PythonModules in SerialEM.
3. If the P2 lens was rotated in Microscope Setup, step C-2:
    a. Update the MagnificationTable lines in SerialEMproperties.txt. This saves the new magnification(s) for the square aperture into the SerialEMproperties.txt file so that it can be accessed for data collection.
        i. In SerialEM: (Taskbar) Calibrations > List Mags
        ii. Compare the listed magnifications with the MagnificationTable in SerialEMproperties.txt. Update the MagnificationTable with any new lines and changes.
    b. Perform a stage shift calibration at the lowest square beam magnification: Perform a stage shift calibration at the lowest square beam magnification in a contiguous series: https://bio3d.colorado.edu/SerialEM/hlp/html/menu_calibration.htm#hid_calibration_stageshift
        i. Use cross-grating with latex beads
        ii. Activate whole image correlation
        iii. In SerialEM: (Taskbar) Calibrations > Image & Stage Shift > Whole Image Corr
        iv. In SerialEM: (Taskbar) Calibrations > Image & Stage Shift > Stage Shift
        v. Copy the implied rotation angle to the SerialEMProperties.txt file, replacing the SolvedRotation value.
    c. For each square beam magnification, determine the new pixel sizes and relative rotations. The new pixel size can be determined by using diffraction peaks calibration standards. Insert a cross-grating, then starting from the previously-calibrated magnification, run the next pixel size calibration, then save calibrations. Then, List Relative Rotations and update DeltaRotation and PixelSize in the RotationAndPixel table in SerialEMproperties.txt.
        i. For high magnifications, in SerialEM (Taskbar): File > New Montage > 3x3 or 5x5 montage > Use Stage movement instead of Image shift > Use Record magnification
            A. The absolute rotation value can be calibrated by fiducial alignment of a tilt series.
            B. There is ambiguity between the true angle and the angle 180 degrees away for determination of defocus handedness (this is independent of physical handedness). This can be checked by measuring the defocus gradient across the tilt axis of tilted images. This can be determined automatically in some tomography software, like Warp, AreTomo3, IMOD, and EMAN2.

ii. In SerialEM (Taskbar): Calibration > Pixel Size > Find Pixel Size
iii. If the RotationAndPixel values for the square magnifications are not contiguous with the previous lines in the table, then the absolute rotation must be calibrated for the first value in that list.

d. When the new pixel size has been found,
   i. In SerialEM (Taskbar): Calibration > Pixel Size > List Relative Rotations
   ii. Update the appropriate RotationAndPixel line in SerialEMproperties.txt with the new pixel size and P value.
e. Calibrate ImageShift and BeamShift in SerialEM for the new magnification(s).
   i. In SerialEM: (Taskbar) Calibration > Image & Stage Shift > Image Shift
   ii. Insert a round aperture at the square beam magnification for the BeamShift calibration.
   iii. In SerialEM: (Taskbar) Calibration > Beam & Spot > Beam Shift
   iv. In SerialEM: (Taskbar) Calibrations > Administrator
   v. In SerialEM: (Taskbar) Calibrations > Save calibrations

#### B. PACE-tomo

1. Download the latest untested version of PACE-tomo, which contains the modifications required for square beam data collection.
2. Download PACEtomo_selectTargets.py. Note that this script is no longer actively supported, and the reader is encouraged to use SPACEtomo for target selection instead.
3. Download PACEtomo_measureOffset.py.
4. Transfer these scripts to the computer running SerialEM and read in these scripts into SerialEM.
   a. In SerialEM: (Taskbar) Script > Edit > Edit # (choose an empty script) > Load > Script of interest > Save As / Save.

### Data collection setup

Most of the data collection setup steps for the square beam follow the same procedures as for the round beam. For steps where the entire sensor needs to be illuminated by a beam, the square beam should be spread to cover the entire sensor.

#### A. Microscope alignments and setup over vacuum

| | |
|---|---|
| **i NOTE** | Follow your facility's best practices for aligning your microscope. Exact steps and settings depend on the microscope configuration. Here, we describe the steps done for a Titan Krios G3i equipped with a Gatan K3 camera with an energy filter. |

##### A-1. General SerialEM setup

1. In SerialEM: (Sidebar) Low Dose Control > Check 'Low Dose Mode'
2. In SerialEM: (Sidebar) Low Dose Control > Set up appropriate magnifications for Focus, Trial, Record, View, and Search presets.
3. In SerialEM: If using PACEtomo, (Sidebar) Low Dose Control > 'Define position of area' > for both 'Focus' and 'Trial' > Position on tilt axis: 0.00 µm

##### A-2. Gatan K3 energy filter tuning

1. Move stage to center over some vacuum
2. Set a bright beam at the data collection magnification (typically spot size 2 or 1). Spread the square beam so that the entire sensor is illuminated by the beam.
3. In DigitalMicrograph: (Sidebar) Tune GIF > Center ZLP
4. In DigitalMicrograph: (Sidebar) Tune GIF > Tune GIF > Full Tune

##### A-3. Calibrate electron dose

1. In SerialEM: (Sidebar) Low Dose Control > Go to: Rec.
2. Change spot size to adjust the flux on the K3 to obtain a reasonable value (see Note below).
3. In TUI: Use Direct Alignments > Beam Shift together with a live view on the Gatan K3 to ensure that beam is centered on the sensor.
4. In DigitalMicrograph: Use Ctrl + Left mouse drag to select the illuminated region on the EF-CCD Image to measure flux. Use this to exclude beam edges / unilluminated area of the sensor from dose rate measurement.
5. In SerialEM: Set up Preview preset to use "Half" or "Square", then collect an image. Check that the image does not include any beam edges or unilluminated areas.
6. In SerialEM: (Topbar) Tasks > Calibrate Electron Dose

   a. If electron dose cannot be calibrated because the beam intensity is outside the calibrated range, manually record the flux on the K3 and use the calibrated pixel size to calculate the dose rate:

$$Flux\left(\frac{e}{Å^2\cdot s}\right) = Flux\left(\frac{e}{px\cdot s}\right) \div \left[Pixel\ size\ \left(\frac{Å}{px}\right)\right]^2$$

7. Additional guidance: https://bio3d.colorado.edu/SerialEM/hlp/html/menu_tasks.htm#hid_calibration_electrondose

#### A-4. Set up exposure settings

1. Set up an appropriate exposure time and frame time to obtain the right flux on the sample (see Note below).
2. In SerialEM: (Sidebar) Camera & Script > Setup > Parameter Set 'Record' > Exposure time / Frame time
3. In SerialEM: (Sidebar) Camera & Script > Setup > Parameter Set 'Record' > Acquisition > Single Image
4. In SerialEM: (Sidebar) Camera & Script > Setup > Parameter Set 'Record' > Processing > Unnormalized
   a. Frames can also be saved as Gain Normalized. It will take up more space, but frames can then be cropped without having to crop the gain reference too.
5. In SerialEM: (Sidebar) Camera & Script > Setup > Parameter Set 'Record' > Check 'Save frames'
6. In SerialEM: (Sidebar) Camera & Script > Setup > Parameter Set 'Record' > Set File Options
7. In SerialEM: (Sidebar) Camera & Script > Setup > Parameter Set 'Record' > Set Folder

| | |
|---|---|
| **i NOTE** | A typical total dose on the sample for tomography is 120 e/Å$^2$ per tilt series. |

#### A-5. Collect gain reference

1. In SerialEM: (Sidebar) Low Dose Control > Go to: Rec / Focus / Trial.
2. Spread the beam so that the entire sensor is illuminated by the beam.
3. In DigitalMicrograph: (Taskbar) Camera > Prepare Gain Reference > Collect reference images for Linear Mode > Collect reference images for Counting Mode > Check 'Expert Mode' > Set dose rate to be slightly lower than Record preset flux to account for loss of electrons due to scattering by the sample (see Note in A-3).
   a. Change the spot size and beam intensity to get the appropriate number of counts for Linear and Counting mode.

| | |
|---|---|
| ℹ NOTE | The applicability of gain references to correct images is based on the flux used for calibration and the flux at the detector through the specimen. Thick specimens have more opportunities for electron scattering, which decreases flux at the detector, and samples at 60° tilt will double the apparent thickness. The gain reference calibration flux should be similar to the flux at the detector through the specimen. |

### B. Microscope alignments and setup over cross-grating

#### B-1. Find eucentric height

1. Insert a cross-grating or holey carbon grid onto the stage.
2. Find an intact square for eucentric height determination.
3. In SerialEM: (Taskbar) Tasks > Eucentricity > Rough Eucentricity.
4. In SerialEM: (Taskbar) Focus/Tune > Set Target > -3; (Taskbar) Focus/Tune > Autofocus.
5. Check FFT with (Taskbar) Process > FFT

#### B-2. Direct alignments

1. In SerialEM: (Sidebar) Low Dose Control > Go to: Rec.
2. Ideally, set the defocus of the microscope close to the data collection defocus. This ensures that the beam will be close to the set position under data collection conditions.
3. In TUI: Direct Alignments > Beam tilt pp X / Y
4. In TUI: Direct Alignments > Beam shift, to center beam
5. For the following steps, spread the Record preset beam so that the entire sensor is illuminated by the beam.
6. In SerialEM: (Taskbar) Focus/Tune > Correct Astigmatism by CTF
   a. Not needed if you have a microscope with a Cs corrector.
7. In SerialEM: (Taskbar) Focus/Tune > Coma-free Alignment by CTF
   a. Not needed if you have a microscope with a Cs corrector.
8. In SerialEM: (Taskbar) Focus/Tune > Calibrate Coma vs Image Shift
   a. Not needed if you have a microscope with a Cs corrector.
9. In TUI: Direct Alignments > Beam shift, use MF-X and Y to center while viewing on DigitalMicrograph to center the beam on the camera.

#### B-3. Re-calibrate image shift and beam shift

1. If the microscope has been serviced prior to the current data collection session, or if it has been a long time between data collection sessions on SerialEM, it may be

worthwhile re-running image shift and beam shift calibrations as in SerialEM Setup section A-3-c.

| | |
|---|---|
| **i NOTE** | If the image shift calibrations are off, the tiles of the montage will be skewed with respect to each other. If the beam shift calibrations are off, the beam will not land in the middle of the sensor during large beam-image shifts. |

#### B-4. Align image shift

1. Find a feature that can be recognized across several magnifications and center it in highest magnification with stage movement.
2. In SerialEM: (Sidebar) Camera & Script > Record, to record a high mag image
3. In SerialEM: (Sidebar) Camera & Script > View, to record a medium mag image
4. In SerialEM: Right mouse drag to center the feature in View mag. Use PAGE DOWN and PAGE UP buttons to scroll between image in buffers B and A, to ensure that A is centered the same as B
5. In SerialEM: (Sidebar) Low Dose Control > Shift offsets > 'Set' for View
6. Repeat if necessary
7. In SerialEM: (Sidebar) Camera & Script > Search, to record a low mag image
8. In SerialEM: Right mouse drag to center the feature in Search mag. Use PAGE DOWN and PAGE UP buttons to scroll between image in buffers B and A, to ensure that A is centered the same as B
9. In SerialEM: (Sidebar) Low Dose Control > Shift offsets > 'Set' for Search

### C. PACE-tomo

| | |
|---|---|
| **i NOTE** | Always follow the latest and most updated protocols from the developer at https://github.com/eisfabian/PACEtomo. |

The following PACE-tomo setup steps were adapted from this tutorial:
https://www.youtube.com/playlist?list=PLsDH3b-_SN9M7QIQ1ErWUeqPu2mIzO158

#### C-1. Measure offset

1. Find an intact square on cross-grating.
2. Run the measureOffset.py script.
    a. measureOffset settings:
        i. increment = 5
        ii. maxTilt = 15
        iii. offset = 5
        iv. plot = True

3. Total tilt axis offset is accurate to ~0.1 μm. If there is a residual focus slope found in the tilt series after processing the data, this value can be adjusted for future sessions.
4. In SerialEM: (Sidebar) Image Alignment & Focus > Check 'Center image shift on tilt axis'.

#### C-2. Lamella setup

1. Insert your grid.
2. Navigate to your lamella of choice.
3. Find eucentric height.
    a. In SerialEM (Taskbar): Tasks > Eucentric height > Rough eucentric
4. Collect a low magnification montage of your grid.
5. Collect a medium magnification montage of your lamella.
6. Add points on your lamella.
    a. In SerialEM (Navigator): Add Points
    b. Select areas of interest on the lamella
    c. The first point selected will be the tracking target, so choose a target that is of lower interest, but still contains sufficient features and contrast for good tracking throughout the tilt series.
    d. In SerialEM (Navigator): Stop Adding
    e. In SerialEM (Navigator): Select the first point in the group of points
7. Run selectTargets.py.
    a. In SerialEM (Taskbar): Script > Run > selectTargets
    b. selectTargets settings:
        i. targetPattern = False
        ii. alignToP = False
        iii. size = 1
        iv. Set maxTilt according to your collection settings
        v. tgtMontage = True
        vi. tgtMntSize is used to draw the montage dimensions on the View image, but currently does not support rectangular montage dimensions. To estimate the montage area, please measure on the View image directly and estimate the montage area by using the pixel size and the expected montage dimensions.
        vii. tgtMntOverlap is also used for montage dimension drawing.
    c. Run the selectTargets script.
        i. Currently, there is not a way to show the full montage region on the preview window of selectTargets. First, decide the area you want to

collect by measuring on the lamella. Then, approximate the montage area and set the montage dimensions and overlap fraction accordingly.

    ii. Choose a directory for saving targets and tilt series.
    iii. Provide a rootname for the PACE-tomo collection area.
    iv. Center the first point at medium magnification with the right mouse button > Press the ‘b’ key to finish.
    v. The script will take a Preview image. If the Preview image needs to be recentered, center the point with the right mouse button > Press the ‘b’ key to finish.
    vi. Repeat until feature is centered to your satisfaction.
    vii. Save the image and coordinates > Yes
    viii. Continue for all targets.

8. Run PACEtomo.py
    a. PACEtomo settings:
        i. Set tomography and data collection settings as desired.
        ii. tgtMontage = True
        iii. Set tgtMntSize to cover your area of interest.
        iv. Set tgtMntOverlap to a minimum of 5%, ideally 10-15%.
        v. tgtMntFocusCor = True
    b. If you have only one Acquire target in the Navigator, select it, in SerialEM (Taskbar): Script > Run > PACEtomo
    c. If you have multiple Acquire targets in the Navigator, select the first one, then in SerialEM (Taskbar): Navigator > Acquire at Items
        i. Select the PACEtomo script
        ii. Set an area for ZLP alignment, if needed.

### Data processing

An updated data pre-processing walkthrough can be found here: https://github.com/hamid13r/sqap-montage/blob/main/walkthrough/walk-through.md

#### B. Align and reconstruct the montage volume

1. Use your favorite tomogram alignment and reconstruction program to align and reconstruct the montage tomogram. It can be helpful to work with a smaller montage or use the tracking tilt series for initial alignments and tilt axis refinement, before working on the larger or full montage. We provide an example workflow below.
2. Tilt axis refinement
    a. In this square aperture system by alignment of the P2 lens, the image rotates with beam intensity. As such, the tilt axis rotation value may slightly differ from the calibrated value.
    b. Only the tracking target contains the physical tilt axis, so the tracking target tilt series will be used to refine the tilt axis rotation value through AreTomo2.
    c. In AreTomo2, use the parameter '-TiltAxis [estimate] 1' to search within a range of ±3° from the supplied value.
3. Montage tilt series alignment and tomogram reconstruction
    a. Due to the potentially large size of montage files, care should be taken with optimizing alignments and minimizing tomogram size at high binning factors prior to commitment to low binning factors for downstream processing.
    b. If tilt series include an unmilled edge of the lamella, removing the out-of-plane feature using IMOD ccderaser may improve alignments.
    c. Import tilt series into WarpTools. Reference: https://warpem.github.io/warp/user_guide/warptools/quick_start_warptools_tilt_series/
        i. Organize files according to WarpTools directory structure.
        ii. Create WarpTools settings files
        iii. In the .mrc.mdoc file, replace the .tif extension with .mrc.
        iv. Create a directory "warp_frameseries". Within warp_frameseries, create a directory "average".
        v. Symbolic link montage tilt images into the "average" directory
        vi. Use *fs_ctf* to estimate defocus
        vii. During *ts_import*, use the '--override_axis <angle>' parameter calling the calibrated tilt axis rotation value.
    d. Perform automated tilt series alignment using AreTomo2 through the WarpTools wrapper. Using a '--patches <X>x<Y>' parameter matching the montage tile dimensions may be useful to compensate for minor deviations in montage stitching.

e. Level the specimen within the tomogram using *ts_autolevel*. This removes excessive tilt in the XZ and YZ views, allowing for a smaller Z dimension and data footprint.
f. Reconstruct tomogram with *ts_reconstruct*. Use the '--halfmap_tilts' parameter to reconstruct tomograms from even-odd tilt images for downstream denoising.
g. [Optional] Improve tilt series alignments with MissAlignment https://github.com/warpem/miss-alignment
    i. In the config.yaml file, change the local refinement image warping grid to match montage tile dimensions.
    ii. Note: This can be very time intensive depending on the size of the montage.

4. Denoising and missing wedge correction
    a. Interpretability of montage tomograms may be improved by programs for denoising and missing wedge correction. In this case, we use IsoNet2 using weighted back projection tomograms reconstructed from even-odd tilt images.
    b. IsoNet2 uses subtomograms for training, so the memory requirements to train denoising and refinement are independent of tomogram size. However, there are CPU memory limitations for writing of tomograms.
        i. Large montage tomograms may be split using IMOD trimvol, processed, and re-assembled using IMOD assemblevol.
5. Segmentation
    a. Membrane segmentation is automatically performed using MemBrain-Seg. Processing of large montage tomograms may be VRAM constrained.
        i. Large montage tomograms may be split using IMOD trimvol, processed, and re-assembled using IMOD assemblevol.
